# Dataset on Temperature Dependency of Zebrafish Early Development

**DOI:** 10.1101/2025.01.28.635255

**Authors:** Angelina Miller, Katja Lisa Schröder, Karsten Eike Braun, Caitlin Steindorf, Richard Ottermanns, Martina Roß-Nickoll, Thomas Backhaus

**Affiliations:** Institute for Environmental Research, RWTH Aachen University, Aachen, Germany; Department of Biological and Environmental Sciences, University of Gothenburg, 40530 Gothenburg, Sweden

## Abstract

Zebrafish (*Danio rerio*) early development stages that do not feed independently, are classified as non-protected life stages under EU Directive 2010/63. Zebrafish reach the independently feeding stage not earlier than 120 hours post fertilization, depending on the incubation temperature. This paper presents a dataset documenting zebrafish early development at two commonly used temperatures 26 °C and 28 °C. We recorded onset of heartbeat and hatching as well as body length, eye size, yolk sac consumption, and swim bladder inflation. Additionally, locomotor activity was tracked after 96 and 119 hours post fertilization.

The dataset serves as a baseline for selecting appropriate experimental conditions and optimizing toxicological study designs. They also facilitate the comparison of experimental results that were recorded at different temperatures. Furthermore, the data provide empirical evidence for amending current guidelines for tests with zebrafish embryos, in particular moving away from a rigid 120 hours post fertilization maximum test duration towards a temperature-dependent maximum test duration that is still in line with the aims of the German Animal Welfare Act.

## Background & Summary

Zebrafish (*Danio rerio*) is one of the most frequently used laboratory animals^1^. Especially early, non-free-living and not independently feeding developmental stages are more and more frequently used for investigating the toxicity of chemicals and other stressors, as such stages do not fall under the legal mandate for animal protection as stipulated by EU Directive 2010/63 on the protection of animals used for scientific purposes^2^. The German Federal Ministry for Environment, Nature Conversation and Nuclear Safety concluded that *D. rerio* life stages younger than 120 hours post fertilization (hpf) do not feed independently and also lack sufficient sensation to feel the effects of the experiments they are subjected to^3^. This 120-hour limit was established based on the following criteria that, if taken together, indicate the ability to feed independently^4^: (1) sufficient development of digestive and (2) ingestive tracts, (3) inflation of the swim bladder, (4) swimming ability and (5) largely depleted yolk.

Strähle and colleagues^4^ concluded that most of these criteria are not met at 120 hpf at an incubation temperature of 28.5 °C. They estimated a developmental delay of almost 20 hours if the incubation temperature is lowered to 26 °C, which would allow to prolong the experiments to 140 hours hpf. However, this temperature dependency of zebrafish development is currently not considered by Directive 2010/63.

In this study we present detailed data that document the temperature-dependent development of zebrafish embryos at 26 °C and 28 °C up to 120 hpf. We recorded the onset of heartbeat and hatching, body length, eye size, yolk sac consumption, and swim bladder inflation. A second data set presents data on the swimming behavior of the eleuthero-embryos acquired at two timepoints from a Light-Dark-Transition Test. All endpoints were recorded for each individual embryo in order to allow quantification of the variability between individuals and experimental groups.

The data provide detailed insights into the temperature dependency of zebrafish development, with a focus on the two most commonly used incubation temperatures. They represent baselines for selecting appropriate experimental conditions in view of experimental goals and, in the case of toxicological studies, the properties of the test chemicals. Given their level of detail, the data are also well suited for optimizing the design of toxicological studies (e.g. they allow appropriate power calculations in order to define minimum detectable effects in dependency of replicate numbers and concentration spacing). The data also facilitate the comparison of experimental results that were recorded at different temperatures. Finally, the data provide empirical evidence for amending the guidelines for tests with zebrafish embryos, in particular moving away from a rigid 120 hpf maximum test duration towards a temperature-dependent maximum test duration that is still in line with the aims of the German Animal Welfare Act^5^.

## Methods

### Chemicals

All chemicals used in this study were purchased from Sigma-Aldrich (Munich, Germany): calcium chloride dihydrate (CaCl_2_·2H_2_O, CAS #10035-04-8), magnesium sulfate heptahydrate (MgSO_4_·7H_2_O, CAS #10034-99-8), sodium hydrogen carbonate (NaHCO_3_, CAS #144-55-8), potassium chloride (KCl, CAS #7447-40-7), benzocaine (C_9_H_11_NO_2_, CAS #94-09-7), ethanol (C_2_H_6_O, CAS #64-17-5, purity: > 99%).

### Zebrafish Husbandry and Egg Selection

Adult, wildtype zebrafish (*Danio rerio*) were maintained in 250 L glass aquaria with water at 26 ± 1 °C and a simulated 14 h daylight period. Fish were fed every second day with TetraMin flakes (Tetra GmbH, Melle, Germany), and Nauplius larvae of Artemia spp. (JBL GmbH & Co. KG, Neuhofen, Germany). For spawning, trays with artificial plants as breeding stimulants were placed in the aquaria. A metal net was employed on the trays to prevent the fish from feeding on their own eggs. Mating, spawning and fertilization took place about 30 minutes after the light turned on at 8:00 a.m. Fertilized oocytes are selected using a microscope (Nikon SMZ1500, Tokyo, Japan) according to the OECD criteria for the fish embryo test^6^. Selected eggs were placed in petri dishes with fish embryo medium prepared according to ISO standard^7^ at 26 °C or 28 °C, respectively.

The embryos were kept at an incubation temperature of 26.06 ± 0.12 °C in the 26 °C incubator (HettCube 200R, Hettich, Beverly, US) and 27.98 ± 0.04 °C in the 28 °C incubator (Innova 40R, Eppendorf, Hamburg, Germany). All embryos were kept in a 14:10 h light:dark cycle. Temperature was continuously recorded with EBI 20-T1 temperature loggers using the Winlog.basic software (ebro by Xylem Analytics, Washington, US).

### Euthanasia

All experiments were completed before 120 hpf and the embryos were euthanized with 40 g/L benzocaine. As zebrafish embryos and eleuthero-embryos are not considered protected life stages before 120 hpf^4^, no authorization for animal experiments was required under European law^2,5^ and the corresponding ordinance in Germany TierSchVersV^8^.

### High-Resolution Time Series

Embryos at the 4-cell to 8-cell stage (as defined by Kimmel and colleagues^9^) were selected for the experiments. 24 embryos were used for each temperature group (26 °C and 28 °C). Temperature groups were split into two sets of 12 embryos each in order to minimize the time spent out of the incubator during photography. Each set was transferred into a 96-round-well-flat-bottom plate containing 300 µL of embryo medium in each well. Individual embryos of each set of 12 were photographed separately using a digital microscope (Keyence VHX-970F, Neu-Isenburg, Germany). The well plate was then immediately closed with the plate lid and placed back in the incubator. The 28 °C plate was sealed with Parafilm (Bemis, Neenah, US) to counteract the higher evaporation caused by the ventilation of the incubator. The order of measurement was: (1) plate 1 at 26 °C, (2) plate 1 at 28 °C, (3) plate 2 at 26 °C, (4) plate 2 at 28 °C.

Imaging was repeated every hour until all 24 embryos had hatched (∼ 68hpf for the 26 °C group, ∼ 60 hpf for the 28 °C group). The developmental status was then documented every two hours until 119 hpf. The time at which the heartbeat began, and the time of hatching were recorded to the nearest hour. The onset of the heartbeat was defined as the timepoint at which first visible contractions of the heart muscle was observed. Additionally, the start of eye and body pigmentation was documented. Body length and eye size were quantified using the microscope software’s internal scale. Fig. 1 shows how the eleuthero-embryo’s body length was measured from head to tail. Due to the refraction of light by the concave water surface in the round well, measured values had to be adjusted to reflect the actual length. A linear calibration model was fitted based on the correlation of objects of the same size with and without water in the well (R^2^ = 0.999).

**Fig. 1:**
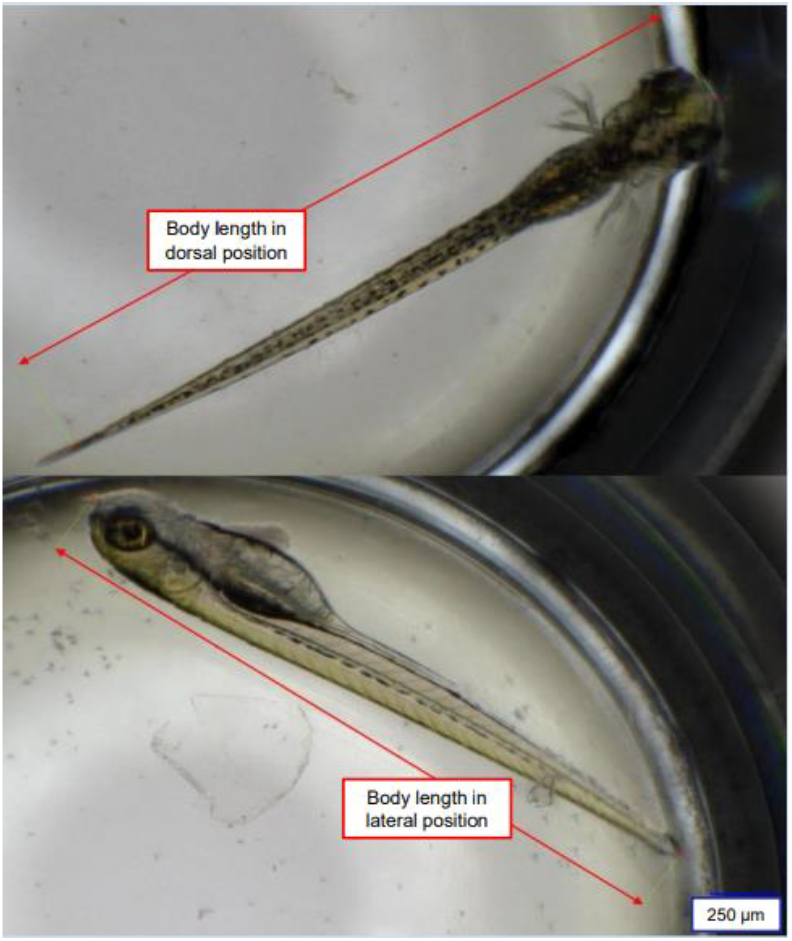
Measurement of zebrafish eleuthero-embryo`s body length in dorsal and lateral views using a 96-round-well-flat bottom-plate. The top image illustrates the dorsal view, with the entire body length measured as indicated by the red line extending from the anterior (head) to the posterior (tail) of the eleuthero-embryo. The bottom image displays the analogue measurement for an eleuthero-embryo in the lateral view. Measurements were obtained using the internal scale of a digital microscope, allowing for precise and consistent data collection.

The whole experiment was repeated twice, resulting in data from a total number of 48 individuals per temperature group.

### Low-Resolution Time Series

72 fertilized eggs (4-cell and 8-cell stages) were selected for each temperature group (26 °C and 28 °C). Otherwise, the experimental conditions were identical to those used in the high-resolution time series with the exception that a single 96-well plate was used for each temperature. After 72 hpf, 96 hpf and 119 hpf, 24 embryos of each temperature group were randomly selected and anaesthetized in 0.4 g benzocaine /L H_2_O for at least ten minutes. The embryos were then individually adjusted with thin, flexible plastic needles, to lie on their sides within a gelatinous fixative (1,67 g dry gelatine /50 mL H_2_O), so that their eyes overlap in the microscopic image (Fig. 2). This allows the comparison of the embryo’s measurements between individuals. In addition to body length and the area of the eye, yolk sac and swim bladder were quantified using the microscope software’s internal scale (Fig. 2). The whole experiment was repeated three times, resulting in data from a total number of 72 individuals per temperature and time group.

**Fig. 2:**
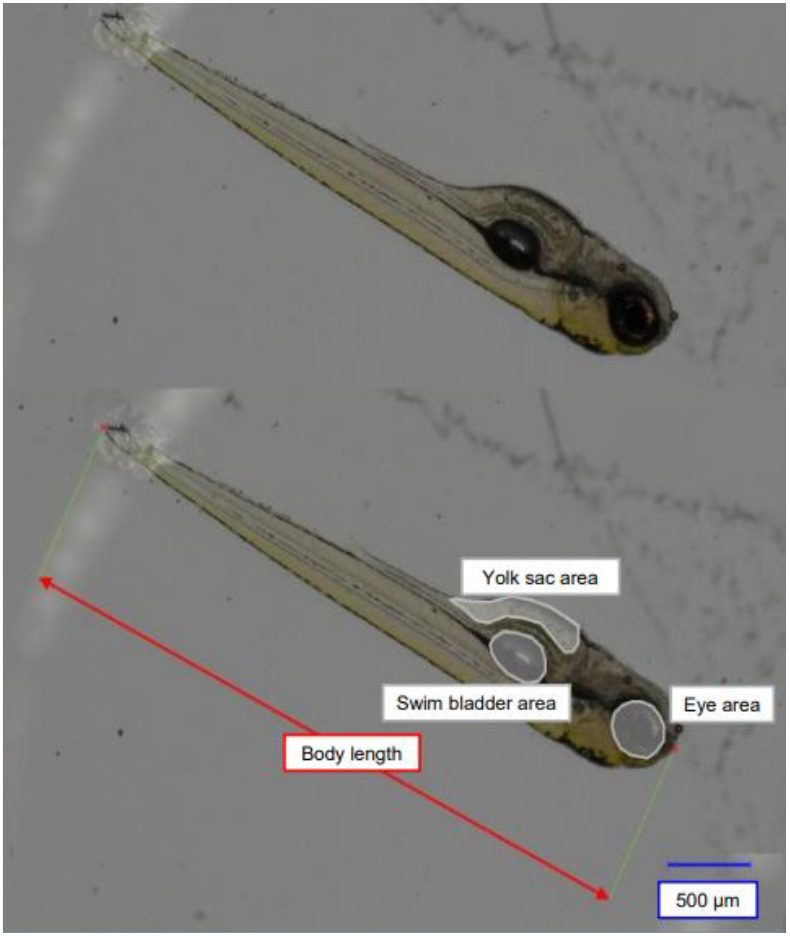
Measurement of body length, eye area, swim bladder area and yolk sac area in zebrafish eleuthero-embryos fixated laterally in gelatine. Measurements were obtained using the internal scale of a digital microscope, providing accurate assessment of key developmental features in zebrafish eleuthero-embryos.

### Light-Dark-Transition Test

For the Light-Dark-Transition Test 96 *D. rerio* embryos from 4 egg batches were raised in 400 µL of embryo medium in 96-well plates at 26 °C and 28 °C, respectively. At 96 hpf and 118 hpf (7:00 a.m., before the onset of light in the incubator) they were exposed in a DanioVision Observation Chamber coupled with a Temperature Control Unit (Noldus Information Technology, Wageningen, Netherlands, at 27 °C), to the following light-dark cycle: (1) 5 minutes light, (2) 5 minutes dark, (3) 10 minutes light and (4) 20 minutes darkness. Phases 1 and 2 served as acclimatization periods. The light intensity was set to 5000 lux. Movement of the embryos was recorded at 25 frames per second. Tracking was performed with EthoVision XT 17.5 software (Noldus Information Technology, Wageningen, Netherlands). Measurements at 96 hpf used eggs from batches 2, 3, and 4, and measurements at 118 hpf used eggs from batches 1, 2, and 3.

## Data Records

All data is publicly available on Zenodo (https://doi.org/10.5281/zenodo.13905257). All data sets are provided as comma separated plain text value files.

### Experiment for High-Resolution Time Series (highres)

Folder *highres* contains the high-resolution time series data for the early developmental stages, heartbeat onset, hatching and body length.

*stages*.*csv* contains one row for each individual embryo in the experiment with columns representing all observed developmental stages. Furthermore, additional columns contain pertinent metadata, such as incubation temperature, replicate number and plate number.

*heartbeat*.*csv* and *hatching*.*csv* include a column indicating the hpf and a column for each temperature which specifies the number embryos that exhibited an observable heartbeat or had hatched at that point in time. An extra column provides the replicate number.

*bodylength_hr*.*csv* has a column indicating the hpf and a column for each well in the plate used for measurements corresponding to one embryo, respectively. The well columns contain the body length of the respective embryo in µm at the associated time point. The columns with metadata provide the associated temperature, replicate and plate number.

### Experiment for Low-Resolution Time Series (lowres)

The low-resolution time series of body length, yolk sack area, eye area, and swim bladder area can be found in the file *lowres*.*csv*. The metadata columns include the following information: hpf, replicate number, and associated temperature.

### Light-Dark-Transition Test (behavior)

Folder *behavior* contains the behavior raw data, which has been divided into two subdirectories by hpf of approximate measurement (*96hpf* and *120hpf*) and again two subdirectories for incubation temperature (*26deg* and *28deg*). The raw data files are designated by a unique batch number. They contain a table with the following columns:

Trial_time [s] - the recorded time point at every 0.04 seconds.

Distance_moved [mm] - the swim distance of the embryo in the respective 0.04 seconds.

Distance_to_point [mm] - the distance kept to the center point of the well.

Turn_angle [deg] - the relative angle the embryo turned during the movement.

Individuum - the location in the well plate as a number from left to right and top to bottom in the plate (A1 = 1, H12 = 96).

Temperature – incubation temperature.

Unit – unit of temperature.

hpf – hours post fertilization.

ID - an individual ID of the embryo in the form w_x_y_z with w = the well position, x = incubation temperature, y = hpf and z = batch number.

## Technical Validation

Temperature loggers were placed in the incubators, in order to validate temperature consistency. The resulting temperature time-series can be seen in Fig. 3 - 5. There were no major spikes in incubation temperatures that were deviating more than 0.2 °C from the nominal temperature. An overall mean between 26.1 °C and 25.9 °C or 28.0 °C and 27.8 °C, respectively was maintained with standard deviations between 0.01 °C and 0.15 °C (26 °C) or 0.02 °C and 0.08 °C (28 °C).

**Fig. 3:**
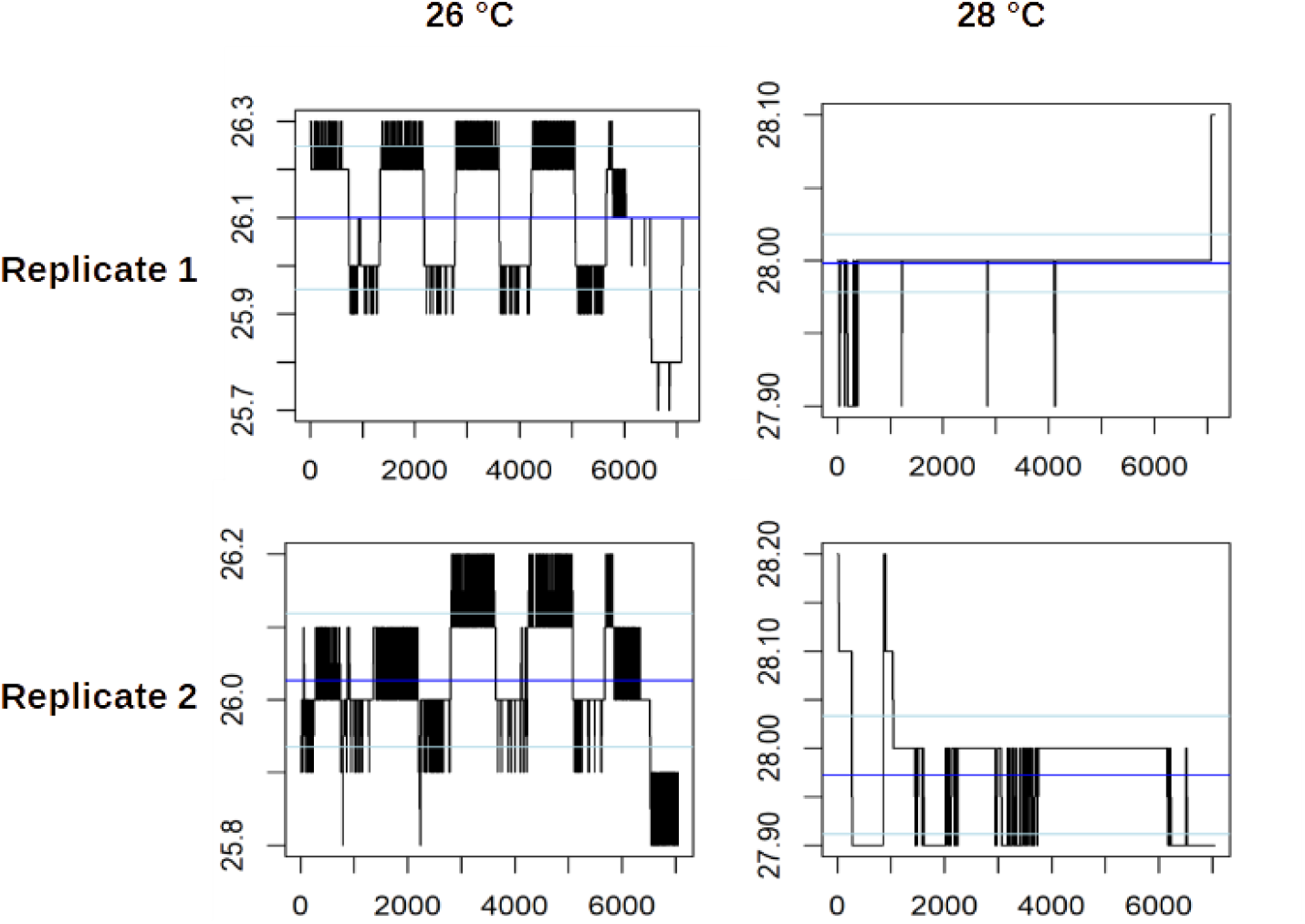
Temperature stability profiles over the time of development for zebrafish eleuthero-embryos used in the High-Resolution Time Series. Incubator temperatures were recorded every 5 minutes for zebrafish eleuthero-embryos incubated at 26 °C (left) and 28 °C (right), with two independent replicates shown. Each plot visualizes temperature fluctuations over time, with blue horizontal lines representing the mean temperature and light blue lines indicating the standard deviations. The y-axis denotes temperature in degrees Celsius, while the x-axis corresponds to the number of recorded data points. Variations in the black bars illustrate the magnitude and frequency of temperature deviations during incubation.

**Fig. 4:**
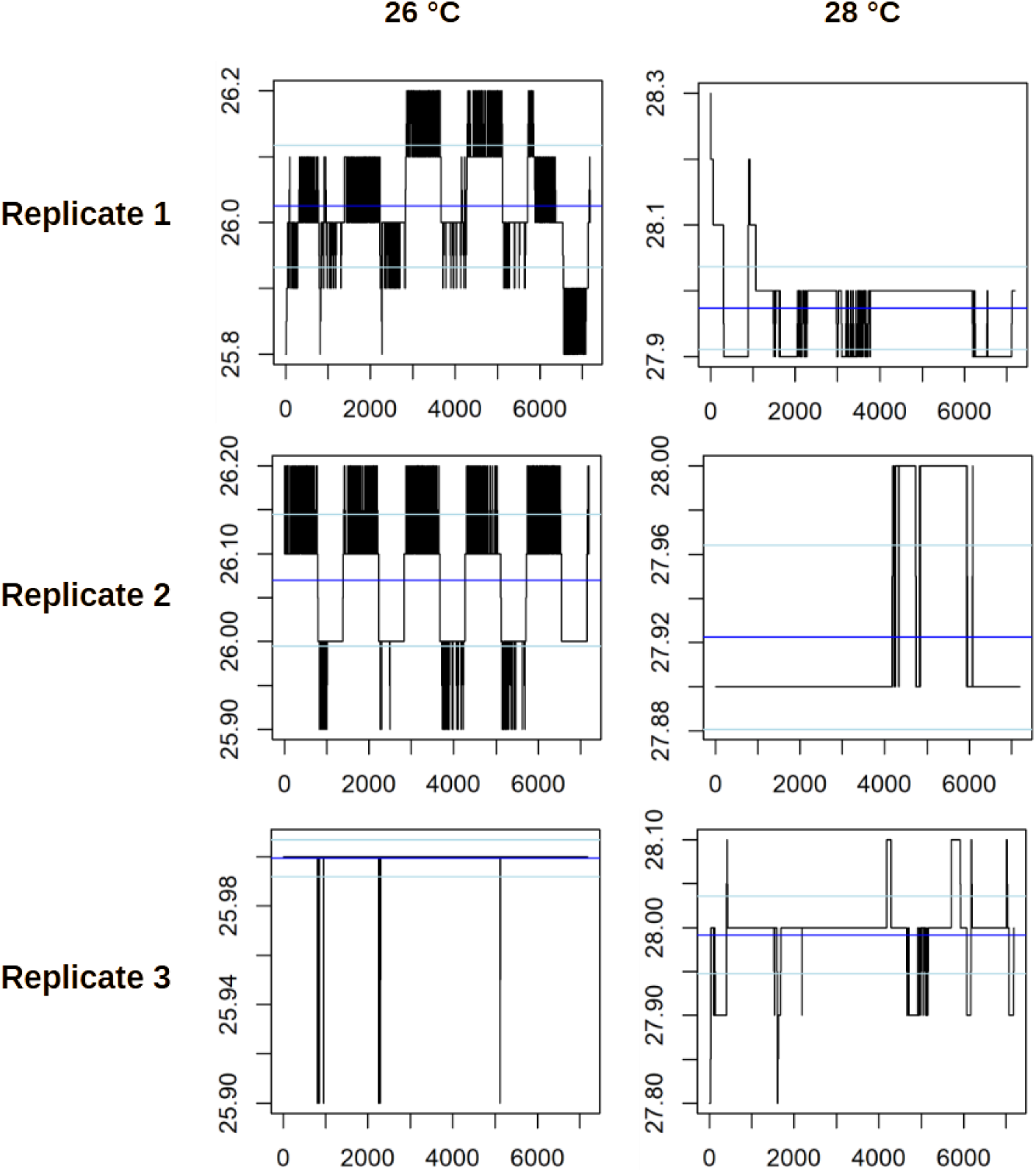
Temperature stability profiles over the time of development for zebrafish eleuthero-embryos used in the Low-Resolution Time Series. Incubator temperatures were recorded every 5 minutes for zebrafish eleuthero-embryos incubated at 26 °C (left) and 28 °C (right), with three independent replicates shown. Each plot visualizes temperature fluctuations over time, with blue horizontal lines representing the mean temperature and light blue lines indicating the standard deviations. The y-axis denotes temperature in degrees Celsius, while the x-axis corresponds to the number of recorded data points. Variations in the black bars illustrate the magnitude and frequency of temperature deviations during incubation.

**Fig. 5:**
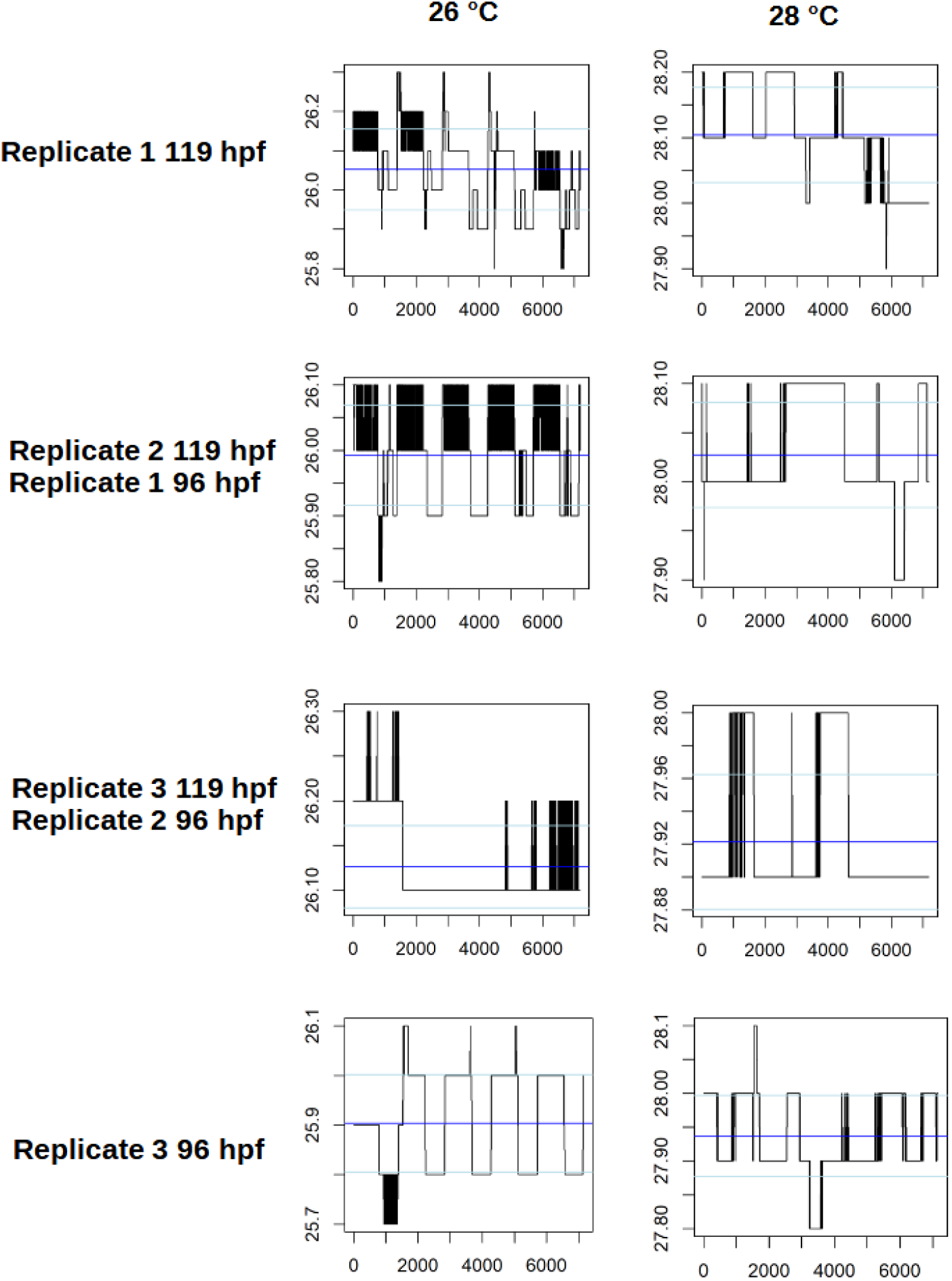
Temperature stability profiles over the time of development for zebrafish eleuthero-embryos used in the Light-Dark-Transition Test. Incubator temperatures were recorded every 5 minutes for zebrafish eleuthero-embryos incubated at 26 °C (left) and 28 °C (right), with three independent replicates for each developmental stage (96 hpf/119 hpf). Each plot visualizes temperature fluctuations over time, with blue horizontal lines representing the mean temperature and light blue lines indicating the standard deviations. The y-axis denotes temperature in degrees Celsius, while the x-axis corresponds to the number of recorded data points. Variations in the black bars illustrate the magnitude and frequency of temperature deviations during incubation.

## Code Availability

The custom code used to assemble the data sets from raw data was developed in R version 4.3.3^10^. It is publicly available on Zenodo (https://doi.org/10.5281/zenodo.13905257) in the repository folder *code* as a single script, named *import_script*.*R*. The script is organized into three sections: highres, lowres, and behavior. Each section begins with comments describing the structure of the raw data, followed by the corresponding processing code for each data set.

## Acknowledgements

This work was funded by the Deutsche Forschungsgemeinschaft (DFG, German Research Foundation) under Germany’s Excellence Strategy – Cluster of Excellence 2186 „The Fuel Science Center” – ID: 390919832.

Funded by the Verein des Hygiene-Institut des Ruhrgebiets e.V., Gelsenkirchen, Germany, funding contract “Standardised and automated evaluation of the zebrafish behaviour test for the environmental toxicological assessment of pollutants” (phase 1 and 2).

## Author contributions

Angelina Miller – conceptualization, methodology, data acquisition, processing, visualization, writing original draft.

Katja Lisa Schröder – conceptualization, methodology, data acquisition, curation, visualization, writing original draft.

Karsten Eike Braun – conceptualization, methodology, data acquisition.

Caitlin Steindorf – data acquisition, processing, writing review and editing.

Richard Ottermanns – project administration, supervision.

Martina Roß-Nickoll – project administration, supervision.

Thomas Backhaus – supervision, writing review and editing.

